# Null results from a dimensional study of error-related negativity (ERN) and self-reported psychiatric symptoms

**DOI:** 10.1101/732594

**Authors:** T. X.F. Seow, E. Benoit, C. Dempsey, M. Jennings, A. Maxwell, M. McDonough, C. M. Gillan

**Author notes:** Correspondence: Tricia Seow, School of Psychology, Áras an Phiarsaigh, Trinity College Dublin, Dublin 2, Ireland.

## Abstract

Alterations in error processing are implicated in a range of DSM-defined psychiatric disorders. For instance, obsessive-compulsive disorder (OCD) and generalised anxiety disorder show enhanced electrophysiological responses to errors – i.e. error-related negativity (ERN) – while others like schizophrenia have an attenuated ERN. However, as diagnostic categories in psychiatry are heterogeneous and also highly intercorrelated, the precise mapping of ERN enhancements and impairments is unclear. To address this, we recorded electroencephalograms (EEG) from 196 participants who performed the Flanker task and collected scores on 9 questionnaires assessing psychiatric symptoms to test if a dimensional framework could reveal specific transdiagnostic clinical manifestations of error processing dysfunctions. Contrary to our hypothesis, we found no association between ERN amplitude and symptom severity of OCD, trait anxiety, depression, social anxiety, impulsivity, eating disorders, alcohol addiction, schizotypy or apathy. A transdiagnostic approach did nothing to improve signal; there was no association between three transdiagnostic dimensions (anxious-depression, compulsive behaviour and intrusive thought and social withdrawal) and ERN magnitude. In these same individuals, we replicated a previously published transdiagnostic association between goal-directed learning and compulsive behaviour and intrusive thought. Associations between the ERN and psychopathology might be smaller than previously assumed and/or dependent on a greater level of symptom severity than other transdiagnostic cognitive biomarkers.

## Introduction

Errors are a critically important information source. They allow us to monitor and continually adapt performance to changes in the environment, to slowly and incrementally improve skills, and to avoid large mistakes by having smaller ones attended to. Without this capacity, we might find ourselves repeating unproductive or damaging behaviours. Conversely, a hypersensitive error detection system might keep us from trying new things, from getting out of our comfort zone and experiencing the learning that comes from failure. Since the early nineties, the mainstay of error monitoring research has been the unique neural response to the commission of errors—the error-related negativity (ERN), a negative deflection of the event-related potential that peaks approximately 50-100ms after an error response (Falkenstein et al., 1991; Gehring et al., 1993). Fundamentally, the ERN represents a well-validated and reliable neurophysiological index of error processing (Holroyd and Coles, 2002) with the anterior cingulate cortex posited to be its neural generator (Debener, 2005; Grützmann et al., 2016; Miltner et al., 2003).

Impairments in error monitoring are phenomenologically characteristic of a range of psychiatric disorders (Ullsperger, 2006), and this has been supported by the frequent observation of alterations in the ERN in patient grous (Gillan et al., 2017; Weinberg et al., 2015a). For example, studies have observed diminished ERNs in schizophrenia (Bates et al., 2002; Foti et al., 2012; Minzenberg et al., 2014; Morris et al., 2006; Simmonite et al., 2012), bipolar disorder (Minzenberg et al., 2014; Morsel et al., 2014) and substance use disorder (Franken et al., 2007; Sokhadze et al., 2008), while enhanced ERN amplitudes are consistently seen in obsessive-compulsive disorder (OCD) (Carrasco et al., 2013b; Endrass et al., 2008, 2014; Endrass and Ullsperger, 2014; Klawohn et al., 2014), social anxiety disorder (Endrass et al., 2014) and generalised anxiety disorder (Carrasco et al., 2013b; Weinberg et al., 2010, 2012a, 2015b). Though the precise functional role of the ERN is still highly debated (Alexander and Brown, 2011; Coles et al., 2001; Holroyd et al., 2005; Vidal et al., 2000; Yeung et al., 2004), there are several interpretations of the various ERN abnormalities observed in psychopathology. For diminished ERNs associated with bipolar disorder and schizophrenia, the phenomenon is hypothesised to reflect internal response monitoring deficits posited to underlie the generation of positive schizophrenia symptoms (Frith and Done, 1988; McGrath, 1991). As for disorders with enhanced ERN amplitudes (i.e. OCD, social anxiety and generalised anxiety), one commonality amongst these disorders is that they are fundamentally characterised by high levels of anxiety. Here, the enhanced ERN is thought to reflect an increased sensitivity to errors (Hajcak, 2012; Weinberg et al., 2012b) which may be experienced as highly distressing (Dreisbach and Fischer, 2012; Hajcak and Foti, 2008; Spunt et al., 2012) in anxiety. This is supported by a body of evidence showing exaggerated physiological changes associated with anxiety (e.g. enhanced startle reflex (Hajcak and Foti, 2008; Riesel et al., 2013), heart rate deceleration (Hajcak et al., 2003a, 2004) and skin conductance changes (Hajcak et al., 2003a, 2004)) are linked to larger ERNs. Given that ERN amplitude shifts are so pervasive in psychiatry, it has been suggested and recognised by the Research Domain Criteria Initiative (RDoC) (Insel et al., 2010) that these reflections of altered error processing may be a transdiagnostic phenomenon (Gillan et al., 2017; Meyer and Klein, 2018; Weinberg et al., 2015a) that holds potential as a biomarker of mental health.

In recent years, meta-analyses have proposed phenotypes beyond Diagnostic and Statistical Manual of Mental Disorders (DSM) categories that may underlie alterations in ERN amplitude and explain their ubiquity across psychiatric groups. Particularly for the enhanced ERN, anxious apprehension (Moser et al., 2013) or uncertainty (Cavanagh and Shackman, 2015) are key candidates, supported by studies in non-clinical samples demonstrating that increased levels of worry (Hajcak et al., 2003b; Moser et al., 2012; Zambrano-Vazquez and Allen, 2014) and threat sensitivity (Weinberg et al., 2016) are associated with larger ERNs. That said, group-level effects in OCD patients tend to be more robust than in generalised anxiety disorder (Endrass and Ullsperger, 2014; Santesso et al., 2006), with a recent meta-analysis positing a higher effect size for OCD than anxiety (Pasion and Barbosa, 2019). As only a few studies have attempted to disentangle (and control for) intercorrelated symptoms within individuals in the same sample, and even fewer have done this in a sample of sufficient size, it remains to be seen if enhancements in the ERN confer risk for anxiety, compulsive symptoms or both.

To test this, we used a dimensional approach whereby we measured co-occurring symptoms of a range of disorders within the same individuals and tested for associations with the ERN in their original form, as well as after they had been reduced to three dimensions—anxious-depression, compulsive behaviour and intrusive thought (hereafter ‘compulsivity’) and social withdrawal (Gillan et al., 2016). Using this method, we previously showed that a transdiagnostic compulsive dimension maps onto deficits in goal-directed control better than OCD symptoms (Gillan et al., 2016); a finding that has since been replicated (Patzelt et al., 2019). We also showed that this method can reveal associations that are hidden by categorical disorder groupings. For example, anxious-depression is linked to reduced confidence, while individuals high on the spectrum of compulsivity have elevated confidence (Rouault et al., 2018; Seow and Gillan, 2019). This finding might explain why group level effects in OCD (where patients have high levels of both compulsivity and anxious-depression) have not revealed confidence abnormalities (Hauser et al., 2017; Vaghi et al., 2017).

Following this methodology, we characterised participants in terms of a broad range of psychopathology (9 questionnaires in total) that have almost all been linked to the ERN in prior work; alcohol addiction, apathy, depression, eating disorders, impulsivity, OCD, schizotypy, social anxiety and trait anxiety. We hypothesised that an enhanced ERN would be associated with OCD, social anxiety and trait anxiety, but that this would be explained by a psychiatric dimension encapsulating high levels of anxiety, i.e. anxious-depression. While we expected OCD symptom severity to correlate with the ERN, we anticipated that the compulsivity dimension would not show an association as prior work has shown diminished ERN in addiction and schizophrenia (see review (Gillan et al., 2017)), both of which are strong contributors to the compulsivity dimension.

We related ERN amplitude to self-report psychiatric symptoms from 196 participants who completed the arrow-version of the Eriksen Flanker task (Eriksen and Eriksen, 1974). Contrary to our hypothesis, we found that none of the psychiatric symptoms nor the transdiagnostic dimensions were significantly associated to alterations in ERN amplitude. To contextualise the absence of ERN affects in the present sample (i.e. effect size), we report results from an additional cognitive task relating goal-directed learning (Daw et al., 2011) to dimensional phenotypes. Here, we did find evidence for an association; replicating prior work (Gillan et al., 2016) showing that goal-directed learning was related to the compulsive behaviour and intrusive thought dimension.

## Results

Participants (N = 196) from a majority student sample completed an arrow-version of the Flanker task, a short IQ evaluation and a battery of self-report questionnaires assessing a range of psychiatric symptoms (see Methods). Individual item-level responses on these questionnaires were transformed into scores for three transdiagnostic dimensions using weights defined in a prior study (Gillan et al., 2016); anxious-depression, compulsive behaviours and intrusive thought and social withdrawal.

### Behavioural results

Across participants, mean error rates ranged from 1.97% to 38.24% (mean (M) = 11.55%, standard deviation (SD) = 7.64%) and mean response times (RT) ranged from 123.73ms to 472.55ms (M = 275.70ms, SD = 67.59ms). We observed basic behavioural patterns expected of the task. Mean error rates increased for incongruent trials (M = 15.58%, SD = 10.08%) relative to congruent trials (M = 3.61%, SD = 4.58%) (*t*_*195*_ = 19.97, 95% Confidence Interval (CI) [0.11, 0.13], *p* < 0.001). Mean RTs were shorter for congruent trials (M = 234.46ms, SD = 66.51ms) versus incongruent trials (M = 296.03ms, SD = 70.16ms) (*t*_*195*_ = −33.38, 95% CI [-0.07, -0.06], *p* < 0.001). Mean RTs were also shorter for error (mean = 212.47ms, SD = 75.20ms) as compared to correct (M = 283.23ms, SD = 66.14ms) trials (*t*_*195*_ = −22.41, 95% CI [-0.08, -0.06], *p* < 0.001). Lastly, post-error mean RTs (M = 294.44ms, SD = 88.47ms) were slower than post-correct mean RT (M = 274.51ms, SD = 68.38ms) (*t*_*195*_ = 6.13, 95% CI [0.01, 0.03], *p* < 0.001). Error rate and RT distributions are visualised in Supplemental Figure S1.

### Response-locked event related potentials (ERPs)

Grand-average ERP waveforms at electrode FCz are presented in Figure 1A for the ERN, correct-related negativity (CRN; negativity succeeding a correct response) and error-related positivity (Pe; positive ERP component succeeding the ERN). ERN and Pe waveforms contained an average of 51.88 (SD = 33.09) error trials per participant while the CRN waveform was constructed with average of 404 (SD = 58.78) correct trials. With the adaptive mean method, the ERN exhibited a mean negative peak of - 3.11μV (SD = 2.79μV) at 37.61ms, CRN a negative peak of 0.30μV (SD = 1.89μV) at 28.59ms, and Pe a positive peak of 3.40μV (SD = 2.54μV) at 192.29ms. Paired t-test indicated more pronounced negativities for the ERN than CRN (*t*_*195*_ = −16.66, 95% CI [-3.82, -3.01], *p* < 0.001) within-subject. Split half-reliability was high for all measures (ERN: *r* = 0.90; CRN: *r* = 0.98; Pe: *r* = 0.90), confirming the suitability of this measure for between-subject analysis.

**Figure 1.**
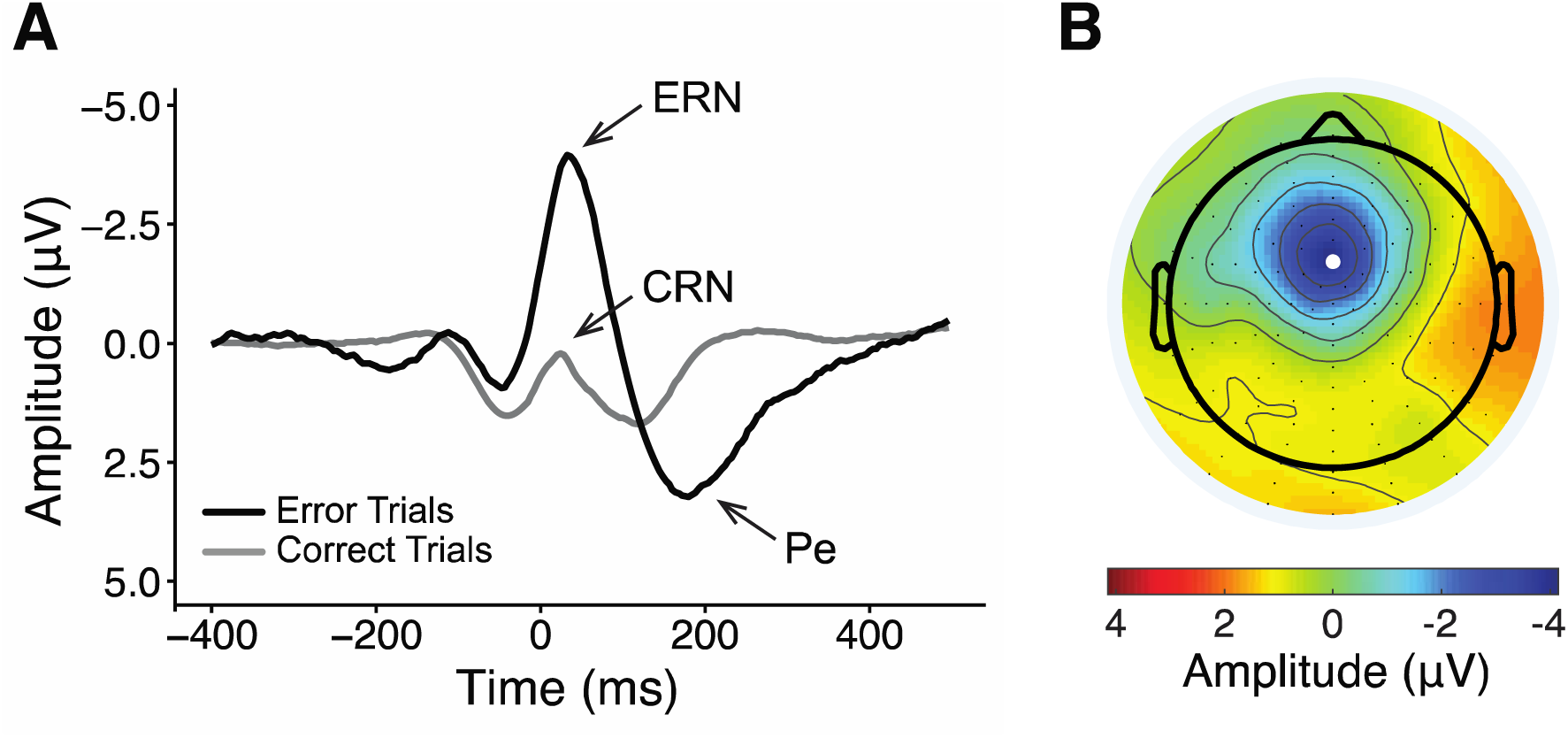
Error-related negativity (ERN). **(A)** Response-locked grand average waveforms for error and correct responses at electrode FCz. Negative values are plotted up. Event-related potential components are labelled: ERN: error-related negativity; CRN: correct-related negativity; Pe: error-related positivity. **(B)** Scalp map displays the voltage distribution at 37.61ms, the average latency of the most negative peak for error trials. Electrode FCz position is indicated with a white dot.

#### Psychiatric symptoms and transdiagnostic dimensions

We tested if ERN amplitudes were associated to the self-reported symptom scores including error rate (*β* = 0.40, *SE* = 0.20, *p* < 0.04) as a control co-variate, which previously have been shown to influence ERN amplitudes (Fischer et al., 2017). In contrast to our hypothesis, none of the psychiatric questionnaires showed a significant relationship to ERN amplitude (all *p* > 0.13, where p < 0.005 is the Bonferroni corrected significance threshold) (Figure 2 and Table 1). For the interested reader, we conducted unplanned, supplementary analyses to test for consistency of our findings across different methods of ERN quantification, electrode site and reaction times (speed-accuracy trade off (Arbel and Donchin, 2009; Gehring et al., 1993)). The patterns of results were remarkably similar: no symptom was significantly related to ERN amplitude in four other ERN quantification methods (all *p* > 0.08, uncorrected) (Supplemental Figure S4) or surrounding electrode sites (all *p* > 0.09, uncorrected) (Supplemental Figure S5), nor influenced by reaction times (all *p* > 0.12, uncorrected). Transdiagnostic phenotyping did not provide a better explanation for the data, with none of the transdiagnostic dimensions significantly associated to ERN amplitude (all *p* > 0.14, uncorrected) (Figure 2 and Table 1).

**Table 1:** Associations between ERN amplitude and total scores of self-report psychiatric questionnaires or transdiagnostic dimensions. SE = standard error. For psychiatric symptoms, each row reflects the (uncorrected for multiple comparisons) results from an independent analysis where each psychiatric symptom score was regressed against ERN amplitude, controlled for error rate. For transdiagnostic dimensions, all three dimensions scores were included in the same regression model.

| **Psychiatric Symptom** | **$\beta$ (SE)** | **z-value** | **p-value** |
| --- | --- | --- | --- |
| Alcohol Addiction | 0.24 (0.20) | 1.23 | 0.22 |
| Apathy | 0.15 (0.20) | 0.75 | 0.46 |
| Depression | 0.15 (0.20) | 0.78 | 0.44 |
| Eating Disorder | 0.08 (0.20) | 0.39 | 0.69 |
| Impulsivity | -0.02 (0.20) | -0.08 | 0.94 |
| OCD | -0.30 (0.20) | -1.52 | 0.13 |
| Schizotypy | -0.11 (0.20) | -0.55 | 0.58 |
| Social Anxiety | -0.13 (0.20) | -0.64 | 0.52 |
| Trait Anxiety | -0.05 (0.20) | -0.24 | 0.81 |
| **Transdiagnostic Dimension** | **$\beta$ (SE)** | **t-value** | **p-value** |
| Anxious-depression | 0.20 (0.22) | 0.91 | 0.37 |
| Compulsive behaviour and intrusive thought | -0.13 (0.22) | -0.57 | 0.57 |
| Social withdrawal | -0.18 (0.23) | -0.81 | 0.42 |

**Figure 2.**
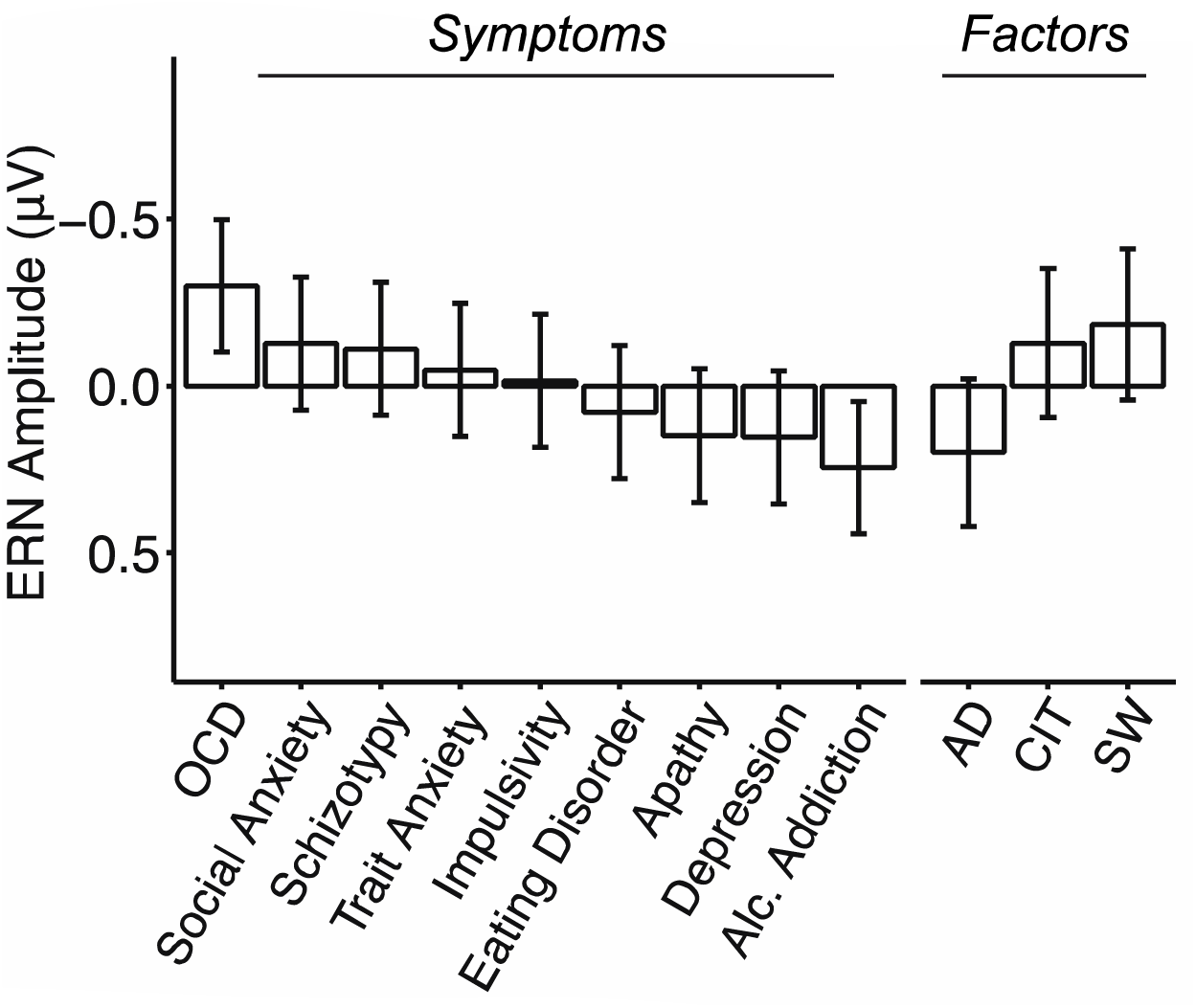
No association between ERN amplitude and self-reported psychopathology. Associations between ERN amplitude with questionnaire total scores or transdiagnostic dimension scores (anxious-depression (AD), compulsive behaviour and intrusive thought (CIT) and social withdrawal (SW)), controlled for error rate. Error bars denote standard errors. Each psychiatric symptom was examined in a separate regression, whereas factors were included in the same model. The Y-axis indicates the change in ERN amplitude as a function of 1 standard deviation (SD) increase of symptom or dimension scores. See Table 1.

### Goal-directed control and compulsivity

For comparison purposes, we also assessed goal-directed learning in the same sample using the two-step reinforcement learning task (Daw et al., 2011). Split half-reliability was *r* = 0.78 for this measure. In prior work, the compulsivity dimension was associated with reduced goal-directed learning (Gillan et al., 2016). We replicated this finding (β = −0.08, *SE* = 0.04, *p* = 0.04) (Figure 3), suggesting that the psychiatric scores obtained from this general population sample were valid, providing a comparator for interpreting the effect size of ERN trends in the present work.

**Figure 3.**
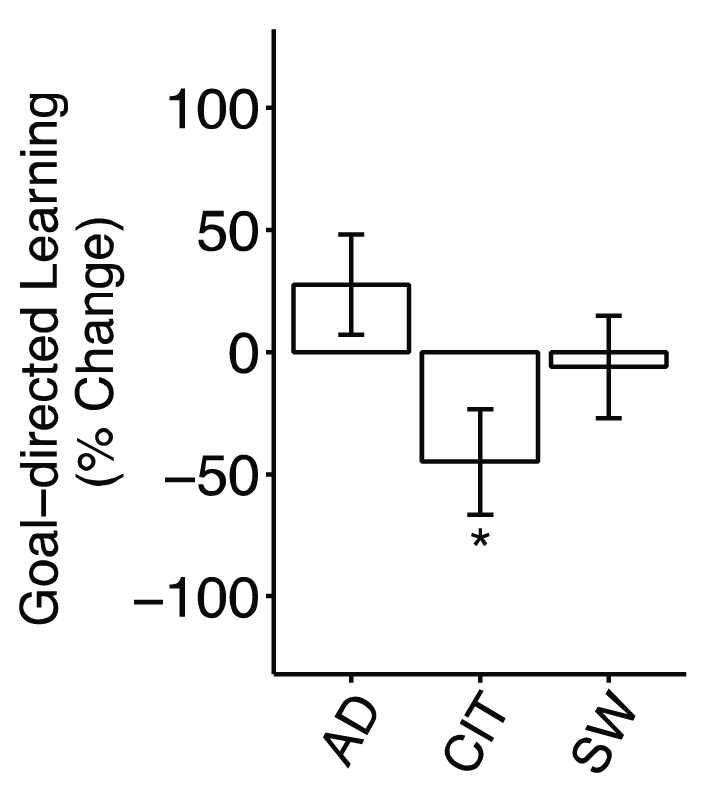
Associations between goal-directed learning and psychiatric dimensions (anxious-depression (AD), compulsive behaviour and intrusive thought (CIT) and social withdrawal (SW)) (N = 196). Error bars denote standard errors. Factors were included in the same model that controlled for age, IQ and gender. The Y-axis indicates the percentage change in goal-directed learning as a function of 1 SD increase of dimension scores. *p < 0.05.

## Discussion

In the present paper we investigated if ERN abnormalities commonly observed in a range of psychiatric disorders could be explained by a transdiagnostic dimension characterised by high levels of anxious-depression. Fundamental to this was the replication of existing associations of ERN amplitude shifts with the clinical phenotypes such as OCD and trait anxiety, but to our surprise we could not detect any significant associations of any symptoms with the ERN. Reformulating questionnaires into transdiagnostic dimensions did not improve signal.

We considered several possible explanations for the data. First, that the range of psychopathology sampled was insufficiently high to detect associations with the ERN. We intentionally enriched our sample by including 8 patients from a local anxiety clinic (who were starting group therapy) to protect against this possibility. This was not necessary; the rest of the sample exhibited high rates of psychopathology (Supplemental Figure S3), consistent with the documented characteristics of university students (Auerbach et al., 2018; Bayram and Bilgel, 2008; Evans et al., 2018). Excluding the 8 patients recruited from an anxiety disorder clinic, 37.97% of the sample who were assessed with a standard psychiatric interview (M.I.N.I., see Methods) presently met criteria for at least one disorder. In terms of the range of self-report symptoms, 25.51% (N = 50) scored of ≥21 on the OCI-R and 54.59% (N = 107) scored of >41 on the STAI, the standard clinical threshold for OCD and anxiety for the respective instruments (Ercan et al., 2015; Foa et al., 2002). Second, we ensured that our data, both self-report and electrophysiological, were valid. Perhaps the strongest evidence for the former is our ability to replicate the specific association between compulsive behaviour and intrusive thought scores and goal-directed planning (Gillan et al., 2016). In terms of the ERN itself, we were able to reproduce all of the expected behavioural (Supplemental Figure S1 and S2) and electrophysiological patterns this task was expected to elicit, suggesting that there were no issues with data quality.

Contextualising these data with the broader literature, ERN abnormalities may be more sensitive to the categorical comparison of patients versus controls than dimensional variation in the general population. Several OCD patient studies did not find any correlation with symptom severity and ERN amplitude within patient groups (Carrasco et al., 2013a; Endrass et al., 2008; Riesel, 2019; Riesel et al., 2017, 2014). The ERN remains elevated in OCD despite successful treatment (Hajcak, 2006; Ladouceur et al., 2018; Schrijvers et al., 2009) and elevations are also observed in unaffected first-degree relatives of patients (Carrasco et al., 2013a; Riesel et al., 2019). As such, the ERN has been couched a psychiatric *vulnerability endophenotype* (Riesel, 2019). Nonetheless, our individual differences approach should have been able to pick up these trait effects along the continuum of scores, regardless of the subtleties of state-based fluctuations.

Perhaps the simplest explanation for these data is that ERN associations with psychopathology are smaller than previously assumed. Notably, effects of OCD symptoms trended in the predicted direction, where individuals who scored higher on this questionnaire had a larger ERN. Likewise, the trend was for alcohol addiction to be associated with a blunted ERN, consistent with the previous literature. Recent reviews add support to this conclusion: two meta-analyses noted that overall effect sizes for anxiety or OCD traits for enhanced ERN were relatively small (Cavanagh and Shackman, 2015; Pasion and Barbosa, 2019) and another that assessed the effect size of ERN amplitude shifts in OCD (Riesel, 2019) noted that larger effect sizes were associated with smaller sample publications, suggesting publication bias. In terms of statistical power, our study has one of the highest sample numbers (N = 196) investigating the ERN in psychiatry. Despite the ERN’s link to psychiatry for two decades (Gillan et al., 2017; Olvet and Hajcak, 2009; Riesel, 2019; Weinberg et al., 2015a), there are only five studies with total N > 150 to date (Hanna et al., 2018, 2016; Meyer and Klein, 2018; Riesel et al., 2019; Weinberg et al., 2015b). Our data suggest that in order to have 80% power to detect a significant association between OCD symptoms and the ERN in a future study, a sample size of N = 729 is required.

This year, several authors have highlighted the potential for a transdiagnostic framework at reconciling the broad range of ERN patterns in the literature (Gillan et al., 2017; Pasion and Barbosa, 2019; Riesel, 2019). The present paper is timely, being the first study to apply an expansive and empirically robust transdiagnostic approach that directly addresses the issue of co-occurring symptoms in a large sample. To our surprise, despite being well-powered, we were unable to significantly replicate previously observed associations with various aspects of mental health and a transdiagnostic approach to quantifying mental health in the sample did nothing to remedy that. Future research in this area might agree that even larger samples than previously assumed are needed to delineate robust associations in general population samples. It is perhaps notable, nonetheless, that our transdiagnostic framework did not perform better in relative terms. Returning to our hypothesis, we found no evidence that anxious-depression might be responsible for the commonly observed enhancement of the ERN in anxiety disorders. In fact, the direction of this non-significant effect was in the opposite direction. This finding might reflect the fact that this dimensional framework is not apt to capture variation in the ERN, in contrast to its application to the study of goal-directed planning and metacognition (Gillan et al., 2016; Rouault et al., 2018). If a dimensional structure exists that can explain the common patterns seen across psychiatric disorders, future research should explore alternatives to the framework employed here.

## Materials and Methods

### Power estimation

An appropriate sample size was determined based on a previous study that reported an association of OCI-R scores and enhanced ERN amplitude that approached significance (*r* = 0.32, *p* = 0.06) (Gründler et al., 2009), an effect size suggesting that N = 155 participants were required to achieve 90% power at 0.005 significance.

### Participants

The majority of participants were recruited from the general public through university channels via flyers and online advertisements, and a small number were patients from St. Patrick’s Mental Health hospital. We included these patients to enrich our sample for self-report mental health symptoms. They were all ≥18 years (with an age limit of 65 years) and had no personal/familial history of epilepsy, no personal history of neurological illness/head trauma nor personal history of unexplained fainting. After reading the study information and consent online, participants provided informed consent by clicking the ‘I give my consent’ button. They also gave written consent before the in-laboratory EEG session. They were paid €20 Euro (€10/hr) upon completion of the study. We collected data from N = 234 participants; N = 8 were patients starting group treatment for anxiety from a local clinic and the rest N = 226 were from the general public. Of the total sample, 138 were female (58.97%) with ages ranging from 18 to 65 (mean = 31.42, standard deviation (SD) = 11.48) years. All study procedures were approved by Trinity College Dublin, School of Psychology Research Ethics Committee and St. Patrick’s Mental Health Services Research Ethics Committee.

### Procedure

Before arriving to the lab for testing, participants navigated a webpage to provide informed consent, basic demographic data (age, gender), list any medications they were currently taking for *a mental health issue* (if so, to indicate the name, dosage and duration) and complete a set of 9 self-report psychiatric questionnaires. For a subset of the participants (N = 110, 47%), they completed a short psychiatric interview in-person on the day of testing (Mini International Neuropsychiatric Interview; M.I.N.I.) (Sheehan et al., 1998). During the experimental EEG session, participants completed two tasks: the modified Eriksen flanker task (Eriksen and Eriksen, 1974) and the two-step reinforcement learning task (Daw et al., 2011). The latter task will be analysed and published separately and so the methods are not described in detail, however we report one basic behavioural result from this task to contextualise our ERN results. Once participants completed both tasks, they completed a short IQ evaluation before being debriefed and compensated for their time.

### Exclusion criteria

Several exclusion criteria were applied to ensure data quality. Participants were excluded if they failed any of the following on a rolling basis. (i) Participants whose EEG data were incomplete (N = 4), corrupted (N = 2) or had all error response-locked epochs over four electrode sites examined (see Response-locked ERPs) failing a threshold criteria of ±50μV (N = 5) were excluded. (ii) Participants who missed >20% of trials (n > 96) of the flanker task were excluded (N = 11). (iii) Participants who scored <55% accuracy were excluded (N = 9). (iv) Participants who incorrectly responded to a “catch” question within the questionnaires: “If you are paying attention to these questions, please select ‘A little’ as your answer” were excluded (N = 7). Combining all exclusion criteria, 38 participants (16.24%) were excluded. 196 participants were left for analysis (115 females (58.67%), between 18-65 ages (mean = 30.82, SD = 11.53 years).

### Disorder prevalence (M.I.N.I.)

After exclusion, 87 participants (44.39%) completed the M.I.N.I., which was introduced part-way through the study. Of these participants, 38 (43.68%) presently met the criteria for one or more disorder. Broken down by recruitment arm, 8 (100%) from the clinical arm met criteria, while 30 (37.97%) from university channels met criteria. This rate is close to published reports on the prevalence of mental health disorders in college student samples (Auerbach et al., 2018; Evans et al., 2018). Of the total sample, 31 (15.82%) were currently medicated for a mental health issue. Broken down by recruitment arm, all individuals recruited from the clinic were medicated, while 23 (12.23%) of those recruited through normal channels were medicated. Further diagnostic information of the sample is summarised in Supplemental Table S1.

### Flanker task

Participants completed an arrow-version of the Eriksen Flanker task (Eriksen and Eriksen, 1974). Each trial consisted of either congruent (<<**<**<< or >>**>**>>) or incongruent (<<**>**<< or >>**<**>>) arrow stimuli presented in white on a grey background of a 32 × 24 cm computer monitor. Participants were instructed to respond as quickly and accurately as possible. Flanker stimulus were presented for 200ms and they had 1050ms to respond by pressing one of two keyboard keys in order to identify the direction of the central arrow. Responses were indicated using the left (‘Q’) and right (‘P’) keys. There were a total of 480 trials split into two blocks, each with 240 stimuli (80 congruent, 160 incongruent) presented. At the end of the first block, if participants had >25% missed trials or had accuracy >90%, they were told to ‘Please try to respond faster!’ for the second block. If their accuracy was <75%, they were told ‘Please try to respond more accurately!’. Otherwise they were told ‘Great job!’. Participants completed 30 practice trials (10 congruent; 20 incongruent) of a slower version of the task prior to the beginning of the experimental task (stimulus presentation: 400ms, response time: 1000ms).

### Behavioural data pre-processing

Missed trials were excluded from analysis. A total of 2275 trials (2.42%) were removed (per participant mean = 11.61 trials).

### EEG recording & pre-processing

Scalp voltage was measured using 128 electrodes in a stretch-lycra cap (BioSemi). EEG signals were sampled at 512 Hz. EEG data were processed offline using EEGLab (Delorme and Makeig, 2004) version 14.1.2 in MATLAB R2018a (The MathWorks, Natick, MA). Data were downsampled to 250 Hz and high-pass filtered at 0.5 Hz. Line noise was removed with CleanLine (Mullen, 2012) at frequencies 50, 100, 150, 200 and 250 Hz. Data were further pre-processed with Clean Rawdata plugin: Bad channels were rejected with a criterion of 80% minimum channel correlation and continuous data were corrected using Artifact Subspace Reconstruction (ASR) (Mullen et al., 2013), with correction parameters set at 10 SD for burst criterion and 25% of contaminated channels for time window criterion. All removed channels were interpolated, and the data were re-referenced to the average. ICA was run with *runica*, pca option on. Ocular and other non-EEG artefacts were rejected automatically with Multiple Artifact Rejection Algorithm (MARA) (Winkler et al., 2011) at a threshold of >40% probability.

### Response-locked ERPs

For quantifying error-related negativity (ERN) characteristics, data were epoched as error trials from -400ms to 500ms, baseline adjusted using a -400ms to -200ms pre-response window. Epoches were rejected with a threshold criterion ±50μV. A total of 36 trials (0.35%) were removed (per participant mean = 0.18 epoches). ERN amplitudes were measured at electrode FCz with the adaptive mean method; the largest negative peak was identified for each participant by searching for the largest preceding negativity within -20ms to 120ms post-response, and the amplitude 40ms before and after the peak was extracted and averaged within-participant. Other ERN measures were also obtained (e.g. non-adaptive mean, minimum amplitude, trough to peak, ERN-correct-related negativity (CRN), average activity over electrodes C22, C23 (FCz), C24 and D2) (see Supplement).

### Self-report psychiatric questionnaires & IQ

Participants completed self-report questionnaires assessing: *alcohol addiction* using the Alcohol Use Disorder Identification Test (AUDIT) (Saunders et al., 1993), *apathy* using the Apathy Evaluation Scale (AES) (Marin et al., 1991), depression using the Self-Rating Depression Scale (SDS) (Zung, 1965), *eating disorders* using the Eating Attitudes Test (EAT-26) (Garner et al., 1982), *impulsivity* using the Barratt Impulsivity Scale (BIS-11) (Patton et al., 1995), *obsessive-compulsive disorder* (OCD) using the Obsessive-Compulsive Inventory - Revised (OCI-R) (Foa et al., 2002), *schizotypy* scores using the Short Scales for Measuring Schizotypy (SSMS) (Mason et al., 2005), *social anxiety* using the Liebowitz Social Anxiety Scale (LSAS) (Liebowitz, 1987) and *trait anxiety* using the trait portion of the State-Trait Anxiety Inventory (STAI) (Spielberger, 1983). These self-report assessments were fully randomized within the psychiatric assessment component of the procedure and were chosen specifically to enable transdiagnostic analysis with psychiatric dimensions described in prior work (Gillan et al., 2016; Rouault et al., 2018). A proxy for IQ was also collected using the International Cognitive Ability Resource (I-CAR) (Condon and Revelle, 2014) sample test which includes 4 item types of three-dimensional rotation, letter and number series, matrix reasoning and verbal reasoning (16 items total).

### Transdiagnostic factors

Raw scores of the 209 individual items from the 9 questionnaires were transformed into dimension scores (anxious-depression, compulsive behaviour and intrusive thought (‘compulsivity’), and social withdrawal), based on weights derived from a larger previous study (Gillan et al., 2016) (N = 1413), that has since been replicated (Rouault et al., 2018).

### Split-half reliability

Internal consistency was calculated for ERN, CRN and error-related positivity (Pe) amplitude. Data were split into two subsets (even versus odd trials/epochs), correlated and adjusted with Spearman-Brown prediction formula.

### Linear regressions

Regression analyses were conducted using linear models written in R, version 3.6.0 via RStudio version 1.2.1335 (http://cran.us.r-project.org) with the *lm()* function. We tested for an association of ERN amplitude with error rate as a z-scored regressor with the equation: ERN ∼ Error Rate. We then investigated if psychiatric symptom severity was related to ERN amplitude shifts controlled for error rate by including the total score for each questionnaire (*QuestionnaireScore*; z-scored) as a predictor in the model above. Separate regressions were performed for each individual symptom due to high correlations across the different psychiatric questionnaires. The model was specified as: ERN ∼ Error Rate *+ QuestionnaireScore*. For the transdiagnostic analysis, we included all three factors in the same model, as correlation across variables was lessened in this formulation and thus more interpretable (only 3 moderately correlated variables r = 0.33 to 0.39, instead of 9 that ranged from r = −0.05 to 0.75). We replaced *QuestionnaireScore* in the equation described previously with the three psychiatric dimensions scores (*anxious-depression, compulsivity, social withdrawal;* all z-scored) entered as predictors. The model was: ERN ∼ Error Rate *+ Anxious-depression + Compulsivity + Social withdrawal.*

### Power calculation

Sample size calculation for a future study was calculated with *pwr.r.test* function in R utilising the correlation coefficient from the equation ERN ∼ OCI-R scores.

### Goal-directed learning

Participants also completed a reinforcement learning task (Daw et al., 2011) that enabled individual estimations of goal-directed learning, which has previously been shown to be deficient in high compulsive individuals (Gillan et al., 2016). Briefly, the task consisted of two stages; in the first stage, participants had to choose between two items that had different probabilities of transitioning (rare: 30% or common: 70%) to one of two possible second stages. In the second stage, participants again had to choose between another two items which were associated with a distinct probability of being rewarded that drifted over time. Individuals performing goal-directed learning (‘model-based’ learner) would make decisions based on the history of rewards and the transition structure of the task, as opposed to individuals who disregarded the transition structure and made decisions solely on the history of rewards (‘model-free’ learner). To quantify goal-directed learning, we implemented a logistic regression model testing if participants’ choice behaviour was influenced by the reward, transition and their interaction of the previous trial, controlled for age, gender and IQ. We then tested the relationship of psychiatric dimensions with goal-directed learning by including the three factors (*anxious-depression, compulsivity, social withdrawal*) into the basic model as z-scored predictors. See Supplemental Methods for details of the regression equations.

The code and data to reproduce the ERN analyses of the paper are freely available at https://osf.io/vjda6/.

## Acknowledgements

TXFS and CMG conceived of and designed the study and analysis plan. MM facilitated participant recruitment for the clinical arm. TXFS coded the experiment, collected and analysed the data. EB, CD, MJ and AM collected the data. TXFS, MM and CG wrote the paper.

This work is supported by a Postgraduate Ussher fellowship from Trinity College Dublin to TXFS and a fellowship from MQ: transforming mental health (MQ16IP13) to CMG.

## Declaration of Interests

None.

## Notes

https://osf.io/vjda6/

