## Supplementary Information for "Null results from a dimensional study of error-related negativity (ERN) and self-reported psychiatric symptoms"

for

*Supplemental Methods*

**ERN and demographics.** In existing work, age, gender and IQ (Falkenstein et al., 2001; Fischer et al., 2016; Larson et al., 2016; Zijlmans et al., 2019) yield various relationships with ERN ampltiude. We explored their effects on the ERN in our data. None of them (all *p* > 0.05) were significantly associated to ERN amplitude shifts, nor did inclusion of them change the model testing the effect of symptom scores on ERN amplitude (all *p* > 0.14, uncorrected).

**ERN amplitude measures.** In the literature, there are various ways to quantify ERN amplitude (Clayson et al., 2013). Here we report the supplementary analyses showing that the main results were not due to our chosen analysis approach – whether it was from an ERN quantification method (Supplemental Figure S4, method details below), or electrode site (Supplemental Figure S5).

For non-adaptive mean, ERN amplitude is extracted at 37.61ms post-response, which is the mean latency of the most negative peak across participants. For peak, the most negative peak was identified for each participant by searching for the largest preceding negativity within -20ms to 120ms post-response. For trough-peak, the trough was identified for each participant by searching for the largest preceding positivity within -100ms before the peak, and the amplitude between the trough and peak was extracted. For ERN-CRN, CRN amplitudes were likewise measured at electrode FCz with the adaptive mean method—the most negative peak was identified for each participant by searching for the largest preceding negativity within -20ms to 120ms post-response, and the amplitude 40ms before and after the peak was extracted and averaged within-subject. The difference of the ERN (with adaptive mean method) and CRN amplitudes was then calculated.

**Goal-directed learning.** The same sample of participants (N = 234) completed the two-step reinforcement learning task (Daw et al., 2011). Several exclusion criteria were applied to ensure data quality, on a rolling basis. i) Participants who responded with the same key in stage one >90% (n = 135) of the time (N = 10). ii) Participants whose probability of staying after common, rewarded trials was less than 5% likely to be at chance, based on a binomial distribution with 50% (chance) probability and the total number of common-rewarded trials experienced by each participant (N = 11). iii) Participants who missed >20% (n = 30) of the trials were excluded (N = 3). (iv) Participants who incorrectly responded to a “catch” question within the questionnaires: “If you are paying attention to these questions, please select ‘A little’ as your answer” were excluded (N = 7). (v) As we intend to analyse the EEG data collected for this task, we additionally excluded participants whose EEG data were incomplete (N = 5) or corrupt (N = 2) from the analysis. 38 participants (16.24%) were excluded in total, leaving 196 participants for analysis. To clean the task data, we excluded individual trials with very fast reaction times (<150ms) reflecting inattention or poor responding. Including missed trials, a total of 1114 (3.77%) trials were excluded.

To estimate goal-directed learning, we performed logistic regression via mixed-effects models with the *lme4* package in R, with Bound Optimization by Quadratic Approximation (bobyqa) with 1e5 functional evaluations. The basic model tested if participants’ choice behaviour to *Stay* or switch relative to previous choice (stay: 1, switch: 0) was influenced by the previous trial’s *Reward* (rewarded: 1, unrewarded: -1), *Transition* (common (70%): 1, rare (30%): -1) and their interaction, with age, gender and IQ as z-scored fixed-effects covariates. Within-subject factors (the intercept, main effects of reward, transition, and their interaction) were taken as random effects (i.e. allowed to vary across participants). In syntax of R, the model was: Stay ~ Reward * Transition + (Age + Gender + IQ) + (Reward * Transition + 1 | Subject). The interaction effect between Reward and Transition was significant, indicating a contribution of goal-directed learning to choice behaviour (*β* = 0.16, *SE* = 0.04, *p* < 0.001). To test if symptom dimensions were associated with goal-directed learning deficits, we included the total scores of the three dimensions (anxious-depression, compulsive behaviour and intrusive thought (‘compulsivity’), social withdrawal) as z-scored fixed effect predictors into the basic model described above. The model was: Stay ~ Reward * Transition + (*Anxious-depression + Compulsivity + Social withdrawal* + Age + Gender + IQ) + (Reward * Transition + 1 | Subject). The extent to which a dimension is related to deficits in goal-directed learning was indicated by the presence of a significant Reward*Transition**Dimension* interaction.

*Supplemental Figures and Tables*

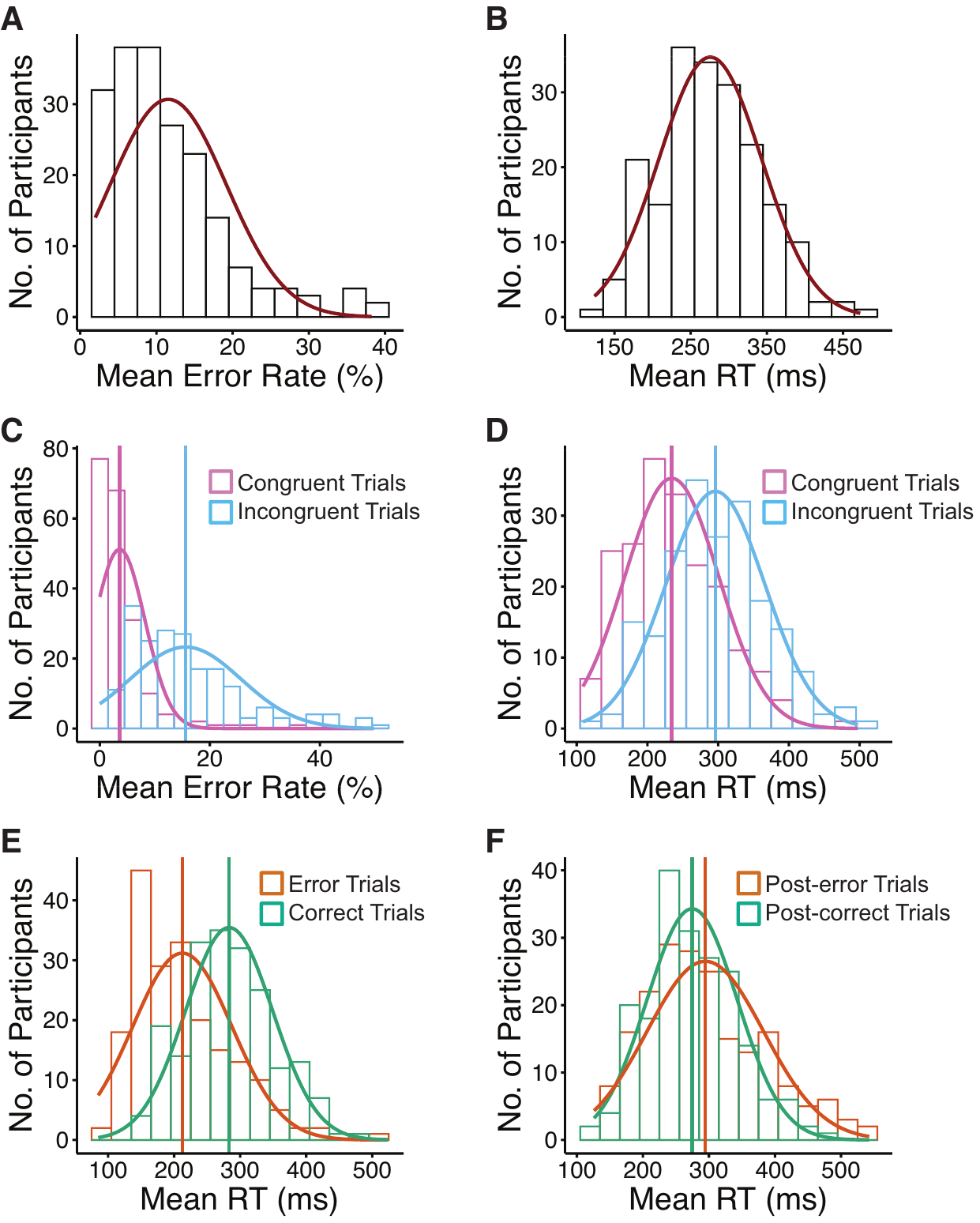

***Figure S1.*** *Across participants, the distribution of:*

***(A)*** *Mean error rate.*

***(B)*** *Mean response time (RT).*

***(C)*** *Mean RT by trial congruency.*

***(D)*** *Mean RT by trial congruency.*

***(E)*** *Mean RT by trial accuracy.*

***(F)*** *Mean RT by post-trial accuracy.*

*Vertical lines denote mean error rate/RT for respective trial type.*

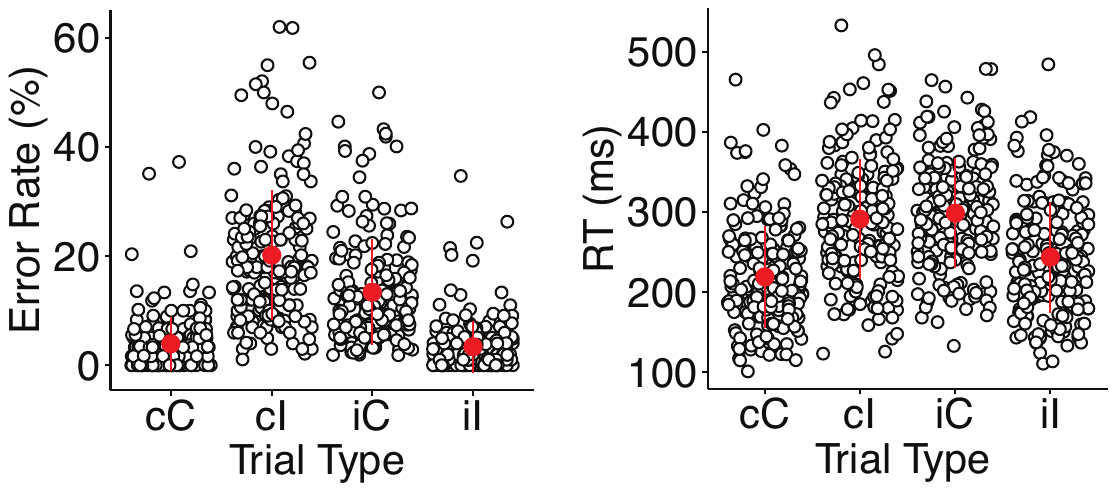

***Figure S2.*** *Mean error rate and response times (RT) for various trial types. cC: congruent trials preceded by a congruent trial, cI: incongruent trial preceded by a congruent trial, iC: congruent trial proceeded by an incongruent trial), iL: incongruent trial preceded by an incongruent trial). White dots represent individual participants, red marker indicates mean and SD.*

**Conflict adaptation.** Conflict adaptation effects refer to the phenomenon wherein previous-trial congruency affects current-trial performance, which have consistently been shown as behavioural adjustment in error rates and RTs in Flanker tasks (Clayson and Larson, 2011; Larson et al., 2016). We replicate these effects, where mean error rates were smaller for iI than for cI trials (*t_195_* = -22.08, 95% CI [-0.18, -0.15], *p* < 0.001) and for cC relative to iC trials (*t_195_* = -15.76, 95% CI [-0.11, -0.08], *p* < 0.001). Additionally, mean RTs were shorter for iI compared to cI (*t_195_* = -24.24, 95% CI (Confidence Interval) [-0.05, -0.04], *p* < 0.001) and for cC relative to iC (*t_195_* = -32.48, 95% CI [-0.08, -0.07], *p* < 0.001) trials.

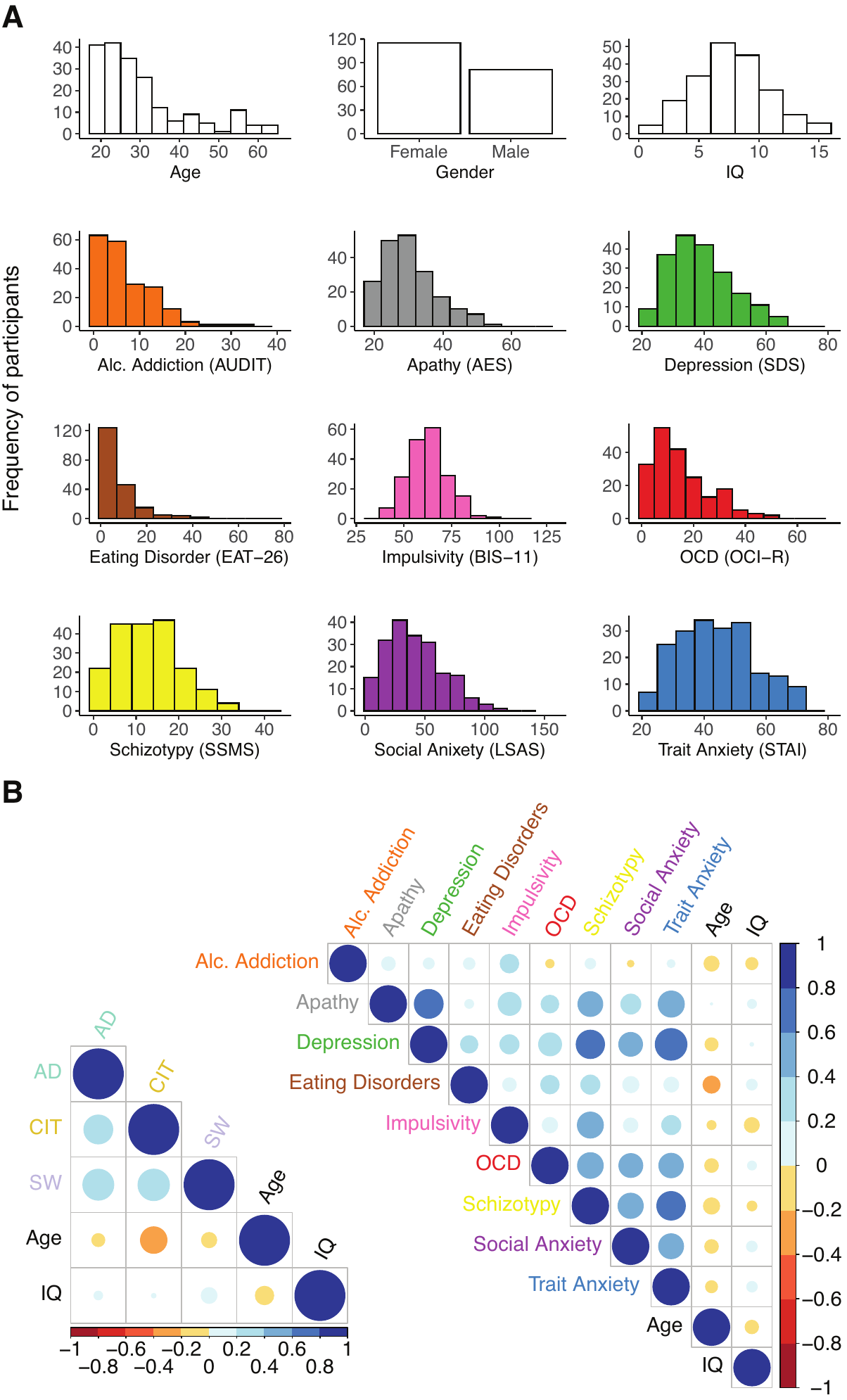

***Figure S3.*** *Demographics and self-reported psychopathology spread.*

***(A)*** *Age, IQ and psychiatric symptoms score distributions across participants.*

***(B)*** *Correlation matrix of mean scores of the nine pyschiatric questionnaires or transformed dimension scores (AD: anxious-depression, CIT: compulsive behaviour and intrusive thought, SW: social withdrawal), including age and IQ. Colour scale indicates correlation coefficient, size of colour patch indicates significance.*

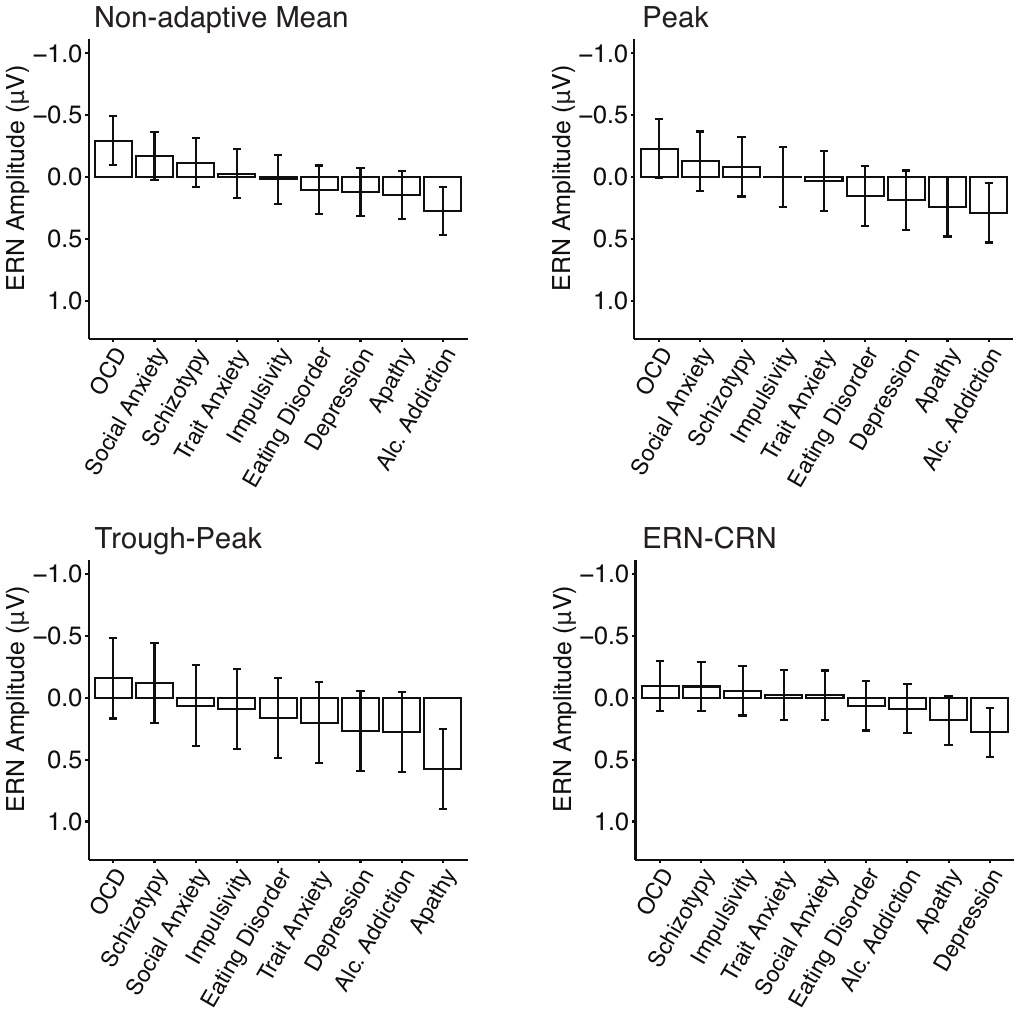

***Figure S4.*** *Associations between psychiatric symptoms with ERN amplitude quantified by various methods at electrode FCz, controlled for error rate. Error bars denote standard errors. The Y-axes indicate the change in ERN amplitude as a function of 1 standard deviation (SD) increase of symptom scores.*

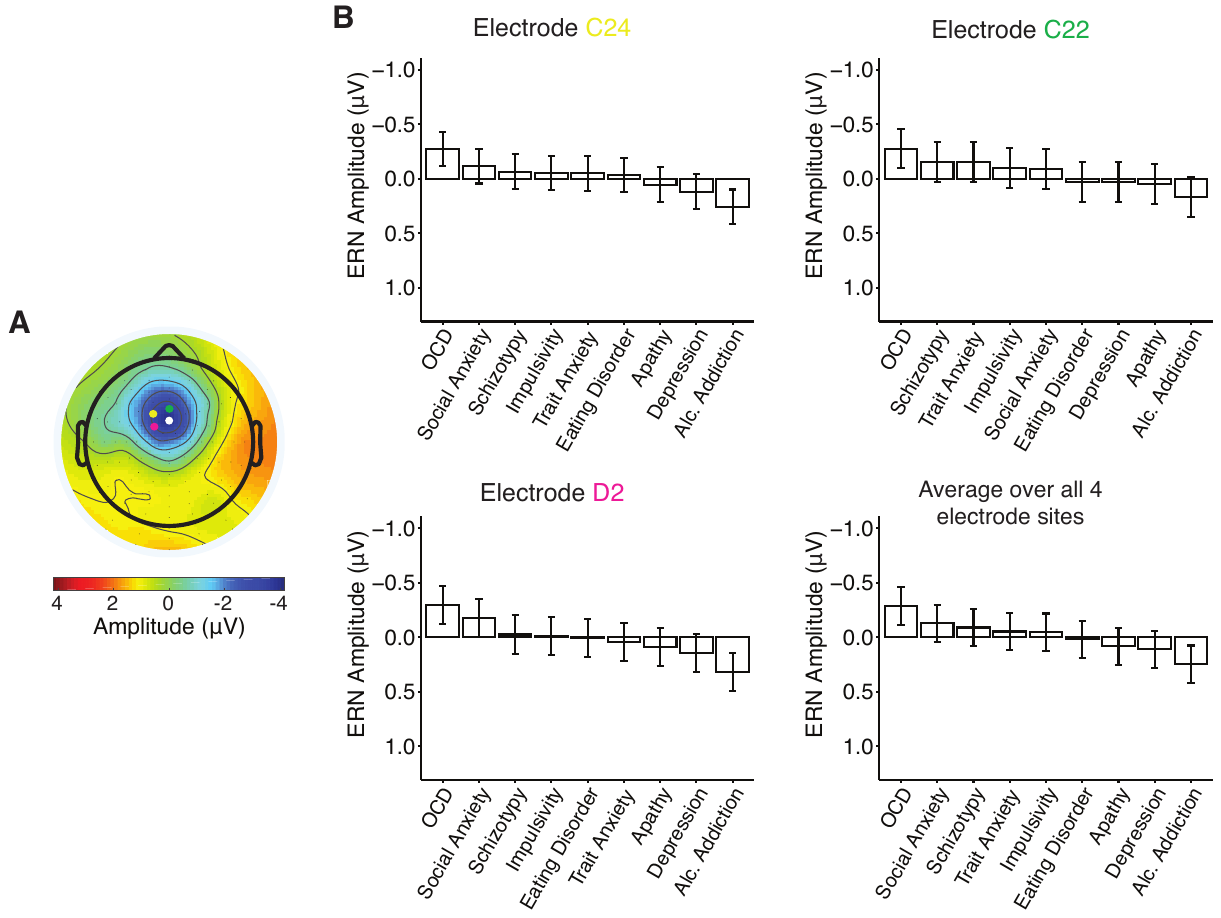

***Figure S5.*** *ERN quantification at various electrode sites with the adaptive mean method.*

***(A)*** *Scalp map displays the voltage distribution at 37.61ms, the average latency of the most negative peak. Coloured dots indicate electrode positions around ERN peak; FCz: white, C22: green, C24: yellow, D2: pink.*

***(B)*** *Associations between psychiatric symptoms with ERN amplitude quantified at various electrode sites, controlled for error rate. Error bars denote standard errors. The Y-axes indicates the change in in ERN amplitude as a function of 1 SD increase of symptom scores.*

***Table S1:*** *Mini International Neuropsychiatric interview (M.I.N.I.) diagnostic information summary for participants who presently met the criteria for at least one DSM-V disorder (N = 38).*

| *Disorder* | *Diagnosis* |
| --- | --- |
| *Mood disorders* | |
| *Major depressive disorder* | *18* |
| *Suicide behavior disorder* | *1* |
| *Bipolar Disorder* | *1* |
| *Anxiety disorders* | |
| *Panic Disorder* | *12* |
| *Agoraphobia* | *4* |
| *Generalised Anxiety Disorder* | *15* |
| *Social Anxiety Disorder* | *11* |
| *Obsessive-Compulsive Disorder* | *5* |
| *Posttraumatic Stress Disorder* | *0* |
| *Substance use disorders* | |
| *Alcohol Use Disorder* | *7* |
| *Substance Use Disorder*  *(Non-alcohol)* | *9* |
| *Psychotic Disorders* | |
| *Psychotic Disorders* | *0* |
| *Eating Disorders* | |
| *Anorexia Nervosa* | *0* |
| *Bulimia Nervosa* | *1* |
| *Binge-Eating Disorder* | *4* |
| *Other disorders* | |
| *Antisocial Personality Disorder* | *1* |
